# Characterization and computational simulation of human Syx, a RhoGEF implicated in glioblastoma

**DOI:** 10.1101/2021.08.26.457821

**Authors:** Ryan J Boyd, Tien L. Olson, James D. Zook, Manuel Aceves, Derek Stein, Wan-Hsin Lin, Felicia M. Craciunescu, Debra T. Hansen, Panos Z. Anastasiadis, Abhishek Singharoy, Petra Fromme

## Abstract

Structural discovery of guanine nucleotide exchange factor (GEF) protein complexes is likely to become increasingly relevant with the development of new therapeutics targeting small GTPases and development of new classes of small molecules that inhibit protein-protein interactions. Syx (also known as PLEKHG5 in humans) is a RhoA GEF implicated in the pathology of glioblastoma (GBM). Here we investigated protein expression and purification of ten different human Syx constructs and performed biophysical characterizations and computational studies that provide insights into why expression of this protein was previously intractable. We show that human Syx can be expressed and isolated and Syx is folded as observed by circular dichroism (CD) spectroscopy and actively binds to RhoA as determined by co-elution during size exclusion chromatography (SEC). This characterization may provide critical insights into the expression and purification of other recalcitrant members of the large class of oncogenic — Diffuse B-cell lymphoma (Dbl) homology GEF proteins. In addition, we performed detailed homology modeling and molecular dynamics simulations on the surface of a physiologically realistic membrane. These simulations reveal novel insights into GEF activity and allosteric modulation by the plekstrin homology (PH) domain. These newly revealed interactions between the GEF PH domain and the membrane embedded region of RhoA support previously unexplained experimental findings regarding the allosteric effects of the PH domain from numerous activity studies of Dbl homology GEF proteins. This work establishes new hypotheses for structural interactivity and allosteric signal modulation in Dbl homology RhoGEFs.

## Introduction

Recent successes designing drugs to inhibit small GTPase based drivers of oncogenesis and advances in modulating protein-protein interactions with structurally guided drug design have inspired renewed interest in characterizing the prolific family of small GTPase activating guanine exchange factor (GEF) proteins with the hope of establishing a new class of drug targets against this expansive class of potential oncogenes(1). These proteins are responsible for modulating a diverse array of cell processes. Diffuse B-cell lymphoma (Dbl) family guanine exchange factors (GEFs) are the largest family of GEFs, containing 71-members out of 82 total RhoGEFs in humans(2). Dbl GEFs facilitate the activation of small GTPases. The mechanistic details of GEF interaction with small GTPases have been reviewed previously(3). The tightly controlled activation and localization of the small GTPase RhoA is directly coupled to stress fiber formation, cell mobility, and proliferation pathways via the opposing effects of Rho Activated Kinase (ROCK) and Diaphanous Homologue (Dia)(4). There is a three-fold higher prevalence of Rho activating GEFs (RhoGEFs) as compared to Rho GTPases (22 members in mammals) indicating that the GEFs are likely regulators of activation specificity for these pathways(5). This is further corroborated by the fact that almost all GEF proteins have been shown to be tightly modulated by numerous mechanisms of inhibition or autoinhibition, suggesting that multiple layers of regulation acting on the GEF are needed for correct conditional flow of these signals within the cell(6, 7). Aberrant activation of these signals can be oncogenic, therefore inhibitors to RhoGEF proteins could be potential cancer therapeutics.

Dbl family RhoGEFs are defined by two tandem domains—the DH-PH domains(8). The 170-190 amino acid DH (Dbl homology) domain facilitates the exchange of guanine nucleotide bound within the small GTPase by structurally manipulating two “finger regions” that encapsulate the nucleotide binding pocket. Simultaneously, many GEFs also affect GDP binding with RhoA by moving a magnesium ion held in complex with Thr-37 and Thr-19 of RhoA out of its binding conformation with the phosphate groups of the RhoA-bound GDP(5) thereby reducing binding interactions. The approximately 120 amino acid pleckstrin homology (PH) domain is often responsible for binding phospho-inositide phosphate (PIP) lipid head groups at the inner leaflet of the cell membrane(9, 10). In some cases, the PH domain allosterically activates the GEF activity of the protein or relieves autoinhibition(11–13)0.2. While the mechanism and overall contribution of PH domain allostery are a matter of continuing study and vary in a protein dependent manner, it is quite clear that in general, GEFs are closely regulated in the cell and often have auto-inhibitory domains or are bound by other proteins to repress their activities when and where they are not intended to be active(14, 15). This spatiotemporal control keeps GEFs from spuriously activating their corresponding small GTPases(16).

It is well established that spurious GEF activity and small GTPase activation are drivers of cell migration, cell proliferation and cancer progression(17, 18). Dachsel et al. 2013 and others revealed that Syx is highly expressed in human glioma cells(19, 20). Experimental depletion of Syx in U87 and U251 glioma cells disrupts cell polarity, resulting in a “fried egg” morphology that does not display chemotactic cell movement that is characteristic of cancer proliferation(21). Notably, the inability of Syx depleted cells to migrate was rescued by the expression of exogenous Syx, but not by a Syx mutant with no GEF activity(19). Additionally, depletion of Syx in conventional or patient-derived GBM cell lines inhibits GBM cell growth (unpublished observations). These results suggest that inhibition of Syx activity may be a possible treatment modality for GBM, and therefore Syx warrants biophysical and structural characterization, to facilitate structurally guided drug design(22, 23). Several structures of Dbl homology RhoGEF DH-PH domains have been solved previously, and GEF characterization methods have been established. In contrast, Syx is in a subgroup of Dbl homology GEFs that are largely uncharacterized outside of basic protein-protein interaction data(24) and no structural information is yet available for this protein(2). Structural elucidation and drug screening efforts require production and purification of milligram quantities of monomeric protein. Failure to overcome protein expression and purification issues are the most common pitfall of structural characterization projects(25).

Here we report the first high yeild expression, and biophysical characterization of purified human RhoGEF Syx including characterization of RhoA binding activity of the Syx DH-PH domain, as well as computational analysis to support ongoing structural studies and drug design efforts.

## Materials and Methods

### Sequence analysis and homology modeling

The full length Syx sequence (UniProt identifier: O94827-1, NCBI reference number: NM_020631.6) was analyzed with the iTASSER homology-modeling server (Table S1)(26) as well as PSIPRED(27), SERp(28), and XtalPred(29) servers. The resultant disorder prediction and structural information were used to guide where truncation would be most appropriate. Several truncation sequences were made based on designing constructs that contained the DH and PH domains but removed unordered regions that would interfere with structural studies. The patterns of truncations were also guided by sequence alignments with Rho GEF structures 1XCG, 1×86, and 3ODO (PDZRhoGEF, LARG, and P115-RhoGEF respectively). Sequences were aligned with the MAFFT server using the L-INS-i method(30).

### Model building and molecular dynamics

Known RhoGEF-RhoA complex structures (1XCG, 1×86, 2RGN, 4XH9, 4DON) were structurally aligned with the Syx homology model to produce an initial Syx-RhoA model. This model was then repeatedly refined using Rosetta docking protocols(31, 32). The resulting homology model of Syx was structurally aligned with several other known structures of PH domains and visually compared to structures containing bound lipid head groups to estimate the orientation of a potential lipid binding pocket on Syx(9, 33–35). The Bio Chemical Library (BCL) software package was used to generate lipid headgroup conformers and Rosetta ligand docking protocols were used to place a PI(4,5)P_2_ lipid into the putative binding pocket(36, 37). The Orientations of Proteins in Membranes (OPM) server and CHARMM-GUI membrane builder were used to add a geranylgeranyl group to the tail of RhoA and then generate an all atom simulated membrane bilayer around the OPM generated lipid-protein interface, as well as place waters and NaCl ions throughout the box(38–40).

Parameter files for GDP, GTP, magnesium, POPC, PI(4,5)P_2_, and geranylgeranyl groups were either generated by CGenFF or found in CHARMM36m params files. NAMD 2.14-CUDA utilizing the CHARMM36m force field was used to run a NPT simulations with 2 fs timesteps for approximately 1.2 microseconds after equilibration(41–43). A temperature of 300° K and 1 atm of pressure was maintained by a Langevin thermostat and barostat and electrostatics were calculated with the particle mesh Ewald method. All simulations were run on GTX1080 Nvidia GPUs until RMSD values reached equilibrium and visual observation confirmed that lipids were correctly oriented(44).

### Dynamic network analysis

Dynamic network analysis was performed as described in Sethi et al. 2009(45). Nodes were defined as C_α_ carbons, or phosphates. Community analysis of groups of residues that are most strongly interconnected was performed using the Girvan-Newman algorithm and visualized with VMD. Optimal path analysis was performed between several residues that clearly bind membrane lipids on both the PH and DH domain of Syx protein and residue Thr-37, located at the center of switch I region of RhoA.

### Protein engineering and mutagenesis

CamSol analysis was performed by uploading the Syx homology model and sequence to the CamSol server(46). The resulting prediction was encoded into the B-factor of the protein and visualized with PyMol. Mutants at these sites were either picked by hand or because they were scored favorably by the Rosetta design protocol(47).

### Expression optimization

Initial expression constructs containing several truncated versions of wild-type mouse Syx homologs were generated by the Anastasiadis lab. Chemo-competent BL21(DE3) cells (New England Biolabs) were transformed according to the manufacturer’s instructions. A single resultant colony was isolated and then used to inoculate a 5 ml liquid culture containing TB containing 12 g/L tryptone, 24 g/L yeast extract, 4 mL/L glycerol, 2.31 g/L KH_2_PO_4_ (17mM), 12.54 g/L K_2_HPO_4_ (72mM), with appropriate antibiotic(48). Cell stocks were made by adding glycerol to 30% and stored at −80°C.

Starter cultures were inoculated by using a sterile pipette tip to transfer a small chunk of frozen glycerol stock into 1ml of pre-warmed TB. After overnight growth, this starter culture was visually checked for cell growth (with a desired OD_600_ of approximately 0.8) and added to an autoclaved 250 ml baffled flask containing 50ml of Terrific Broth (TB) and cells were allowed to grow at 37°C. IPTG was added to a final concentration of 0.5mM when the culture reached an OD_600_ of 0.8, and 1 ml aliquots were taken for analysis at desired time points. For subsequent SDS-PAGE analysis, aliquots were centrifuged at 17K x *g* and the supernatant was discarded. 10µl of cell pellet was mixed with 500µl 1X Laemmli buffer (Bio-Rad Cat# 1610747) and stored at −20°C. Samples were incubated at 95°C for 5 minutes and spun down at 17K x *g* for 10 min to remove cell debris before analysis of raw supernatant was performed via SDS-PAGE using a 12% acrylamide-tris gel and subsequent overnight transfer to a Western blot PVDF membrane and visualization with an anti-His antibody (Qiagen Cat# 34440, RRID:AB_2714179).

Further optimization was done with His6-TEV-Syx_393-792_. This construct was transformed into BL21(DE3)(NEB), BL21(PlysS)(NEB), Lemo21(DE3) (NEB), BL21-AI(Invitrogen), KTD101(DE3), KJ740(DE3), C41(DE3), and C43(DE3) strains of *E. coli* cells. Strain KJ740 was obtained from the Yale *E. coli* Genetic Stock Center (CGSC), and the (DE3) lysogen was made using the λDE3 Lysogenization Kit 538 (EMD Millipore #69734-3). Expression was performed as described above apart from chloramphenicol being used with Lemo strains. Trials with 1 and 2 mM of Rhamnose were tested with the Lemo21(DE3) cells. For BL21-AI, arabinose at 0.2% final concentration was added along with IPTG at induction, and TB medium contained 0.1% glucose.

### Cloning

Codon optimized constructs of the truncated versions of human Syx were designed and obtained from GenScript. Fusion constructs were generated as described previously(48) by cloning our codon optimized Syx gene into vectors containing the following tags that are cleavable with tobacco etch virus (TEV) protease: N-terminal His6 plus maltose binding protein (MBP; RRID:Addgene_29708); C-terminal MBP plus His6 (Addgene_37237); N-terminal His6 plus glutathione S-transferase (GST; RRID:Addgene_29707); N-terminal His6 plus small ubiquitin-like modifier (SUMO; RRID:Addgene_29711); or N-terminal His6 plus green fluorescent protein (GFP; RRID:Addgene_29716). Note that in the plasmid names for this clone the numbering of Syx residues was based on the Syx isoform from NCBI Reference Sequence NP_001036128.1. The sequence of this isoform is identical to the UniProt sequence O94827-1 used for the computational studies, aside from an additional 56 amino acids at the N-terminus of NP_001036128.1. All constructs were transformed into both BL21(DE3) and T7 Express lysY/Iq high competency E. coli (New England Biolabs #C3013I). Ligation independent cloning was performed with the In-Fusion HD Cloning Plus system (Clontech #638910). Plasmid DNA was prepared with the QIAprep Spin Miniprep (QIAGEN #27106). DNA sequences were verified by Sanger sequencing at the DNA Laboratory core facility at Arizona State University or at GenScript.

### Preparation scale E. coli expression

A 5ml overnight growth of the N-terminal His6-MBP-TEV-Syx_393-792_ (referred to as MBP-Syx_393-792_ for brevity, or Syx_393-792_ if referring to protein which has undergone TEV cleavage and MBP removal) construct in T7 Express *lysY/I*^*q*^ E. coli was visually checked for cell growth (OD_600_ of approximately 0.8) before being added to 1 L of pre-warmed TB containing 100ug/ml ampicillin. Cells were grown at 37°C and 300 rpm shaking to an OD_600_ of 0.8. The temperature was decreased to 25°C and IPTG was added to a final concentration of 0.4mM. Cells were allowed to grow for another 4 h before being spun down. Cell pellets were weighed and resuspended in 10ml Lysis Buffer A per 1 g of cells, and the resulting slurry was frozen at −80°C. Lysis Buffer A contained PBS (137 mM NaCl, 2.7mM KCl, 10mM Na_2_PO_4_, 1.8mM KH_2_PO_4_, pH 8), 2mM dithiothretol (DTT), protein inhibitor cocktail (Roche cOmplete Ultra, Sigma part no. 5892791001), 1mM PMSF.

### Purification of E. coli derived MBP-Syx_393-792_

Frozen cells were resuspended in 4°C lysis buffer and 2mg/ml hen egg lysozyme (Sigma part no. 4403) and 0.2mg/ml bovine pancreas DNase (Sigma part no. 9003-98-9) were added. The thawing cell slurry was sonicated on ice at 50% power for 1sec on, 2sec off, for 1min using a Branson 550 sonicator. The resulting slurry was centrifuged at 40,000rcf for 15min at 4°C and then filtered through a 0.45µM filter before using a 150ml superloop connected to an AKTA FPLC in a 4°C cold room to load protein at 0.5ml/min onto a 5ml amylose column (Cytiva Product no. 28918779). The column was previously equilibrated with buffer A (10mM Na_2_PO_4_, 1.8mM KH_2_PO_4_, pH 8, and 1 mM TCEP). The protein was washed with 10 column volumes at 5ml/min and then eluted in 1ml fractions with buffer A plus 50mM maltose. Fractions with an absorbance peak at 280 nm (combined volume of 20-25ml at a protein concentration of 1-5mg/ml) were concentrated to a volume of 500µl using a 30kD Molecular Weight Cut Off (MWCO) spin concentrator (Millipore part no. UFC903096) spun at 3000rcf, while visually ensuring there was no turbidity and mixing the solution every 5min with a pipette. The concentrated sample was injected onto a pre-equilibrated Superdex 200 Increase 30/100 GL column (Cytiva part no. 28990944) and run at 0.4ml/min at 4°C with buffer A. Peaks were pooled and stored at 4°C. The protein concentration and yeild was determined at this stage by absorbance at 280nm(A_280_), using a molar extinction coefficient of 112,355 M^-1^cm^-1^ for the MBP-Syx_393-792_ fusion construct. SDS-PAGE gels were run on all fractions and stained with Coomassie to ascertain purity.

### TEV cleavage and negative purification

The fractions containing purified protein at the expected molecular weight as determined by Coomassie stained SDS-PAGE gel were cleaved with TEV protease by incubating a 10:1 ratio of protein and TEV mixture overnight at 4°C in buffer A. This mix was then incubated with 3ml of nickel NTA slurry for 20min to remove the His tagged MBP and TEV. Flow-through and subsequent washes were collected and concentrated using a 30kDa Molecular Weight Cut Off (MWCO) spin concentrator (Millipore part no. UFC903096). Protein concentrations were confirmed with A_280_ measurements after each interval with a molar extinction coefficient of 39880 M^-1^cm^-1^ for the cleaved Syx_393-792_. Presence of the correct protein species was confirmed routinely with western blot. RhoA (1mg/ml) and anti-Syx antibody were blotted directly on PVDF as control before blocking with BSA (Figure S8). Anti-Syx antibody (Proteintech Cat# 19830-1-AP, RRID:AB_10858324) (8ul in 10ml of TBST) was used in conjunction with a goat anti-rabbit antibody (Jackson ImmunoResearch cat# 111-035-003. RRID: AB_2313567) for visualization.

### Size Exclusion Chromatography

The superdex 200 column was connected to the AKTA FPLC in the 4°C cold room and equilibrated with at least 2 column volumes of sterile filtered water followed by at least 2 column volumes of 25mM HEPES, 150mM NaCl, and 1mM TCEP buffer until the A_280_ and conductance traces appeared constant. Samples were spun down at 17,000g for 10min in a tabletop centrifuge at 4°C before 500µl of sample was injected onto the column and run at 0.4ml/min for the entire run. Fractions were collected at 1.5ml intervals over 1.5 column volumes. After each run, the column was re-equilibrated for 2 column volumes before the next run was initiated. A standard curve was run periodically to determine the elution volume which corresponded to the molecular radius of each protein (Cytiva part no. 28403842). Peaks eluting before 8ml were considered to be in the void volume of this column.

### Dynamic Light Scattering

The monodispersity of the purified protein was ascertained with a Molecular Dimensions SpectroSize 302 DLS apparatus with a 785nm 60mW laser imaging of 2µl protein droplets suspended in a 24 well hanging drop plate. The DLS data were collected in 10 scans with 20min long scans each, and resultant data was examined by the cumulants method^113^. DLS based buffer screening was done by mixing 2ul of protein with 2ul of each well of the Hampton research buffer screen 1 and 2 kits (CAT NO: HR2-072, HR2-413) and incubated for one hour before testing with DLS.

### RhoA expression and purification

RhoA plasmid (RRID:Addgene _73231) expressing the TEV cleavable N-terminal His6-tagged soluble domain of RhoA including residues 1-184 (referred to as His6-TEV-RhoA_1-184_ or just RhoA unless otherwise stated) was transformed into Rosetta 2 BL21(DE3) cells and frozen as glycerol stocks that were used to inoculate 5ml overnight starter cultures grown overnight at 37°C, 250rpm. 5ml overnight starter cultures were used to inoculate 2L baffled flasks containing 1L of TB media with antibiotic and allowed to grow at 37°C until the culture reached an OD_600_ of 0.6–0.8. At this point 250mM of IPTG was added, the temperature was turned down to 18°C, and the culture was allowed to grow overnight. The resulting culture was spun down at 4000 rpm and the pellet weighed and frozen at −80°C.

### RhoA purification

Critically, all buffers were supplemented with 50µM GDP (Sigma cat no. G7127) to maintain RhoA in a folded state. Seven grams of cells were homogenized with 80ml of Lysis Buffer (Tris-HCl pH 8.0, 150mM NaCl, 2mM MgCl_2_, 50*μ*M GDP, 10% glycerol, 5mM imidazole, 1mM PMSF, 1 SIGMAFAST ETDA-free protease inhibitor coctail tab (Sigma sku S8830-20TAB), 2mg/mL lysozyme, 2mM TCEP). Cells were lysed via probe-sonication with a Branson 550 sonicator set to run in intervals of 1 sec on, 2 sec off for a total of 1 min at 50 % power. Sonication was repeated 2 times before the lysate was spun down at 40000rcf for 20min at 4°C. The resulting supernatant was passed through a 0.45μM syringe filter before the supernatant was added to a 150ml superloop attached to a GE Akta series FPLC running unicorn 7 software to automate the following protocol: The clarified supernatant was injected at 1ml/min onto a 5ml Ni-NTA column (Cytiva part no. 17524802) equilibrated with Wash Buffer (50 mM Tris-HCl pH 8.0, 500 mM NaCl, 2 mM, MgCl_2_, 50 *μ*M GDP, 10% glycerol,15 mM imidazole, 2 mM TCEP). The column was washed at 5ml/min with 10 column volumes of wash buffer before a linear gradient of Elution buffer (50 mM Tris-HCl pH 8.0, 300 mM NaCl, 2 mM MgCl_2_, 50 *μ*M GDP,10% Glycerol, 200 mM Imidazole, 2 mM TCEP) was used to elute the bound protein(49).

Fractions of the elution step were collected and run on an SDS-PAGE gel before being stained with Coomassie to reveal fractions containing bands corresponding to the molecular weight of RhoA at 22kDa. Fractions were pooled and an A_280_ absorbance trace was measured using the elution buffer as a blank to estimate protein concentration and overall yield. The presence of His6-TEV-RhoA_1-184_ was also confirmed with Western blot using an anti-RhoA antibody (Thermo Fisher Scientific Cat# MA1-134, RRID:AB_2536840).

### RhoA – MBP-Syx_393-792_ complex formation

Several methods for complex formation were tested to assess the most effective protocol for forming the MBP-Syx_393-792_-RhoA complex to ascertain which produced the highest quality protein. The following protocols used Un-cleaved MBP-Syx_393-792_ (unless otherwise stated) incubated at 4°C with un-cleaved His6-TEV-RhoA_1-184_ in a 1:2 ratio for 1 hour in all 5 trials. In “mix 1” (Figure 3A, purple trace), 10mM of EDTA was added to the mixture to chelate magnesium out of the GDP binding site of RhoA. “mix 2” (Figure 3A, red trace) used cleaved Syx_393-792_ (Figure 3B) mixed with RhoA, and also contained 10mM EDTA to chelate magnesium. “mix 3” (Figure 3, black trace) contained MBP-Syx_393-792_ mixed with RhoA which was buffer exchanged 6 times in a 10k MWCO spin concentrator to remove all buffer containing GDP. In “mix 4” (Figure 3, light blue trace) an excess of ammonium sulfate was used to precipitate a protein mixture containing both MBP-Syx_393-792_ and RhoA in order to to competitively force GDP out of the active site of RhoA and remove GDP containing buffer. For “mix 5” (Figure 3, grey trace) the process was the same as “mix 4” except that the ammonium sulphate precipitation was performed on RhoA alone, then MBP-Syx_393-792_ protein solution was added to the RhoA. Each resulting solution was run on a Superdex 200 column as previously described. Controls show RhoA (Figure 3A, green trace), MBP-Syx_393-792_ (Figure 3A, orange trace), and a molecular weight control (Figure 3A, dark blue trace).

### Circular Dichroism

Cleaved Syx_393-792_ was buffer exchanged 4 times into CD buffer (150 mM sodium fluoride adjusted to pH 7.5 with 50mM monobasic and dibasic sodium phosphate) using a 15 kDa MWCO Amicon spin column. The circular dichroism (CD) spectra were measured on a Jasco J-815 circular dichroism spectrophotometer scanning from 190-260 nm(50). The resulting spectra were analyzed by the servers BeStSeL, K3D2, and DichroWeb’s CDSSTR protocol using the SMP180 basis set(51–53). Spectral analysis was compared to CD spectra of homology models and known protein structures with homology greater than 20% (Table 1) by back-calculating CD spectra with the PDB2CD server to check that the experiment accurately recapitulated secondary structure of the predicted folds seen in the homology model(54).

**Table 1.** Secondary structure prediction by various methodsBack calculated secondary structure prediction of homologous GEF crystal structures generated by PDB2CD are shaded in grey, including a Syx homology model.

| Method | Helix | B-sheet | Turns | Unordered | NRMSD | Homology |
| --- | --- | --- | --- | --- | --- | --- |
| BeStSel | 30.1% | 24.6% | 8.6% | 36.7% | 0.013 |  |
| CDSSTR (SMP180) | 54.0% | 10.0% | 10.0% | 24.0% | 0.014 |  |
| K2D3 | 36.2% | 20.1% |  | 43.8% |  |  |
| Syx model | 47.2% | 10.9% |  |  |  | 100% |
| (PDZRhoGEF) 1XCG | 47.1% | 16.7% |  |  |  | 25.8% |
| (P115-RhoGEF) 3ODW | 57.7% | 14.0% |  |  |  | 23.9% |
| (LARG-RhoGEF) 1X86 | 48.7% | 13.0% |  |  |  | 21.4% |

## Results

### Sequence analyses & homology modeling guided construct optimization for structural studies

To assess which regions of Syx were most likely to be ordered and determine which domains were likely to be useful targets for structural discovery, a homology model and corresponding sequence analysis were performed. Sequence identity between Syx and the most homologous proteins that have solved structures indicated a sequence identity of approximately 25% (Table S1). Homology modeling of Syx DH-PH fragments predictably produced a model largely similar to known homologous structures, however regions of the DH and PH domain have several stretches that were unable to be modeled reliably as determined by their homology to known structures^123^. Most notably a loop on the PH domain had no matching homology anywhere in the PDB (Figure S1). Analyses of the full length Syx protein using iTASSER(55) and DISOPRED3(56) showed that the regions flanking the DH and PH domain were predicted to contain intermittent regions of highly disordered loops and poly-glutamate stretches. These regions were predicted to be poor targets for structural discovery, therefore truncation of the wild type protein was warranted.

### Expression screens generate reliable protein production conditions

HIS-tagged mouse Syx constructs received from the Anastasiadis lab contained Syx GEF domains with truncations over four residue ranges: 1) 290-799, 2) 290-748, 3) 406-799, 4) 406-748, as well as a GST-tagged full-length mouse Syx. Expression in *E. coli* was only observable for mouse construct Syx_406-799_, which showed modest expression with several unwanted lower molecular weight bands for the mouse Syx protein (Figure 1, lane B). Screening of expression conditions revealed that peak protein levels were achieved within four hours of induction at 25°C. At this juncture a human homologue of the Syx_406-799_ mouse construct we refer to as Syx_393-792_ was optimized for expression in *E. coli*. Screening showed meager enhancement of expression, but more importantly it showed none of the unwanted lower molecular weight bands seen in the mouse construct (Figure 1, lane A (human optimized) vs. lane B (mouse native)). Marginal improvements in yield were achieved by using T7 Express *lysY/I*^*q*^ BL21(DE3) *E. coli* (NEB). We then aimed to further improve on the codon-optimized construct with the addition of various fusion proteins for enhanced solubility and purification (Figure S2). We also attempted small scale IMAC (Ion Metal Affinity Chromatography) purification, however, we were not able to recover any substantial amount of protein. When expression was screened on several fusion constructs, the N-terminal MBP-Syx fusion construct produced large quantities of protein which were clearly visible on a Coomassie stained SDS-PAGE gel at the correct molecular weight (Figure S2).

**Figure 1.**
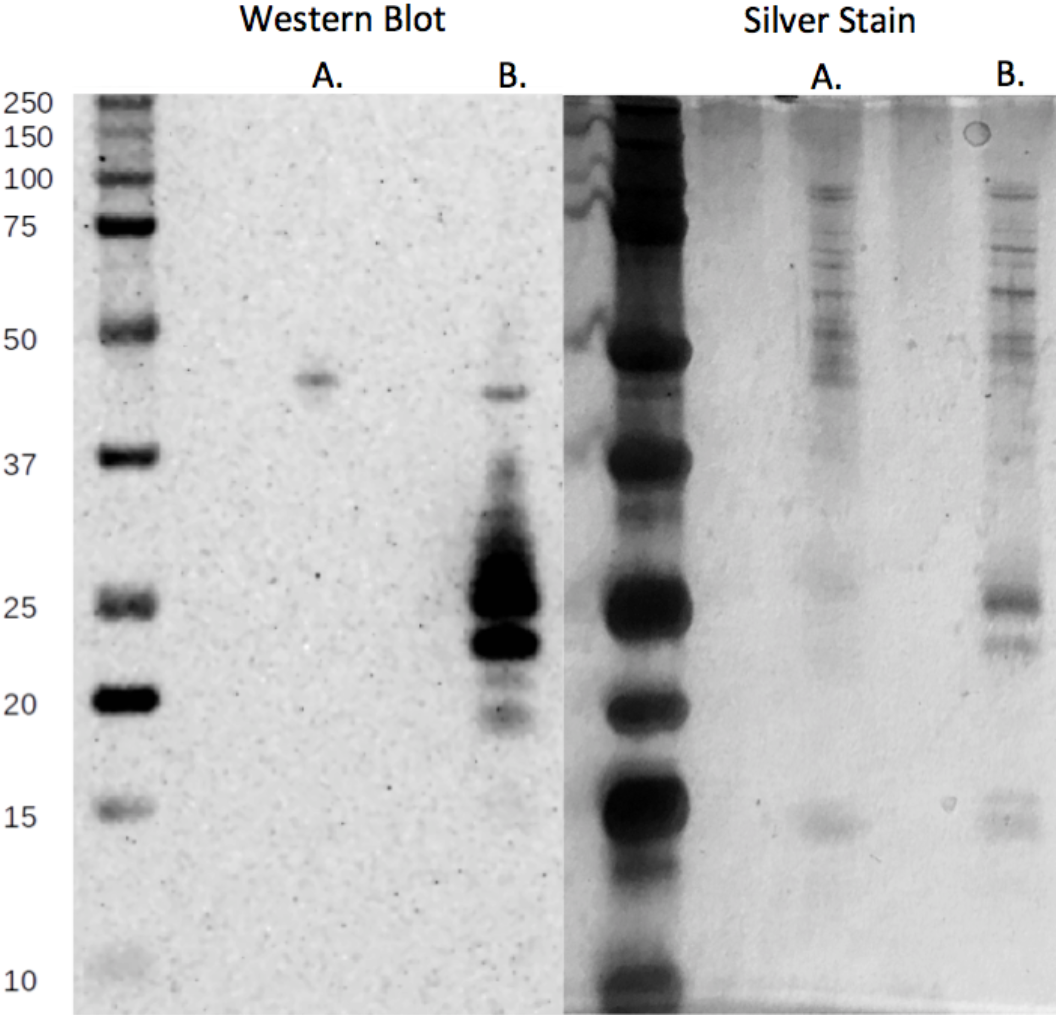
Comparison of expression of mouse Syx_406-799_ and codon optimized human Syx_393-792_ genes. Identical fractions were visualized with silver stain and western blot. (Lanes A. Human Syx_393-792_: MW 48.3kDa. Lanes B. Mouse Syx_406-799_: MW 46.6kD

### Size exclusion chromatography & dynamic light scattering reveal pure, milligram quantities of MBP-Syx_393-792_ with a high molecular radius

Protein quality after expression was tested via size exclusion chromatography (SEC) and DLS. Purification of the MBP-Syx_393-792_ via amylose column and SEC was confirmed to be over 90% pure by Coomassie stained SDS-PAGE (Figure S2). Protein yields after amylose column were approximately 10 mg per liter of culture. DLS showed presence of an aggregate with a particle sized ~17 nm and a polydispersity index (PDI) of 0.54 indicating a mixture of populations (Figure S8). All SEC runs with MBP-Syx_393-792_ on a Superdex-200 increase 30/100 GL SEC column resulted in a large clearly visible peak eluting at or near the void volume of the column, around 8ml (Figure 2). This occurred regardless of changes in pH from pH 10 to pH 5, below which no protein was visible indicating that it had crashed out of solution. In addition, trials of numerous common buffer additives commonly used to reduce aggregation were not effective.

**Figure 2.**
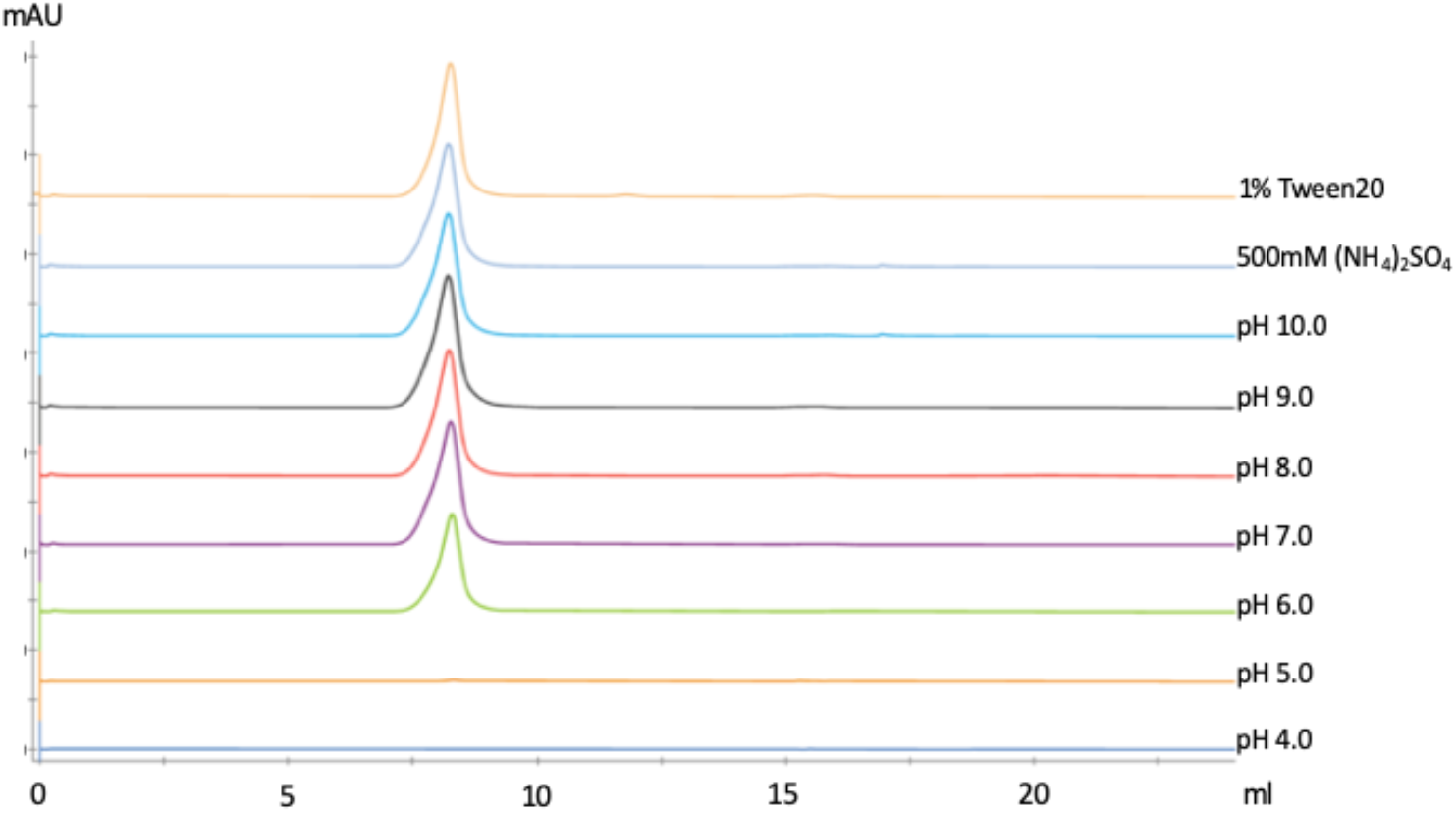
Screening of SEC buffer conditions did not produce monomeric MBP-Syx_393-792_ as shown by SEC absorbance trace at 280nm. Each sample was from the same preperation of MBP-Syx_393-792_ and differed only by the buffer additives indicated in the legend. Buffers all contained 500mM NaCl, and also contained the following buffers: pH 4-5: 20mM sodium acetate buffer, pH 6: 20mM sodium citrate, pH 7: 20mM sodium phosphate, pH 8: 20mM Tris, pH 9: 20mM glycine, and pH 10: 20mM CAPS buffers. 1% Tween20 and 500mM (NH_4_)_2_SO_4_ respectively were both added to a base buffer of 20mM HEPES pH 7.5, 500mM NaCl in the case of the final two SEC runs.

These results are indicative of a soluble aggregate or very large homo-oligomer. The best conditions established (10mM Na_2_PO_4_, 1.8mM KH_2_PO_4_, pH 8) were able to generate a very small peak corresponding to monomeric protein (MBP-Syx_393-792_ control shown in orange Figure 3A). When the peak corresponding to the molecular radius of monomeric protein was isolated and re-run on the same SEC column, the result was a similar equilibrium of two peaks with the predominant population in the void volume. Solubility screening using Hampton detergent and additive screens were not successful at reducing the presence of the aggregate as measured by DLS.

**Figure 3.**
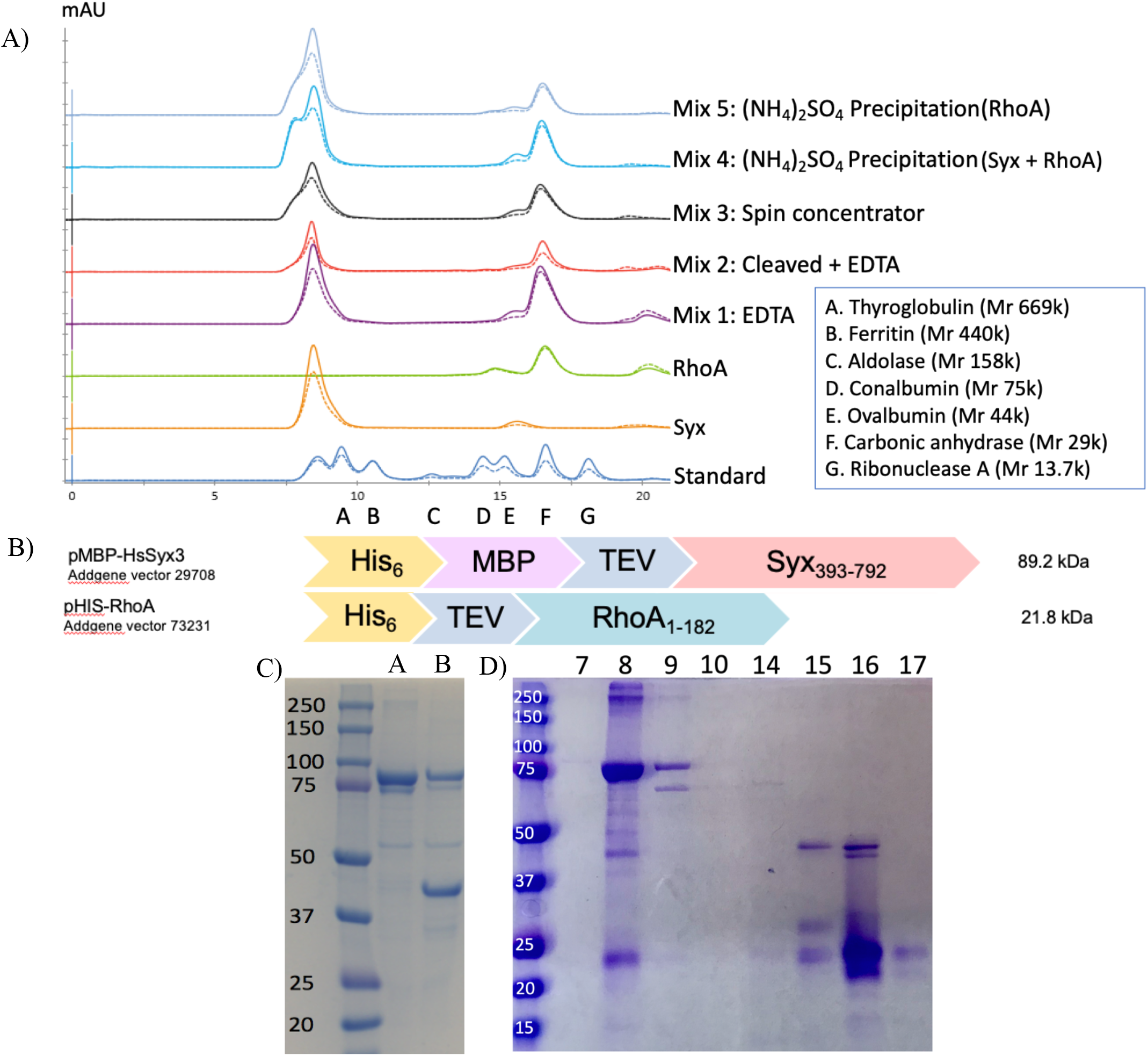
(A) SEC absorbance trace at 280nm of mixtures of MBP-Syx_393-792_ and RhoA after using varying strategies for removing GDP from RhoA to induce the formation of a high affinity MBP-Syx_393-792_–RhoA complex. The protein standard consists of A. Thyroglobulin (Mr 669 000), B. Ferritin (Mr 440 000), C. Aldolase (Mr 158 000), D. Conalbumin (Mr 75 000), E. Ovalbumin (Mr 44 000), F. Carbonic anhydrase (Mr 29 000), G. Ribonuclease A (Mr 13 700). MW of N-term MBP-Syx is 90.8kDa. RhoA is 22kDa. (B) Cartoons depict expressed protein fusion constructs of both MBP-Syx_393-792_ and RhoA. RhoA is truncated at residue 184 to remove its highly charged linker and geranylgeranyl transferase recognition site. Mix 5 was made by ammonium sulphate precipitation of just RhoA before mixing. All other complex formation mixes (Mix 1-4) were made by first mixing MBP-Syx_393-792_ and RhoA 1:1 and allowed to equilibrate for one hour before attempting to remove GDP. (C) MBP-Syx_393-792_ (lane A) cleavage with TEV protease produces an approximately 48kDa band (Lane B) corresponding to the molecular weight of Syx_393-792_ (48kDa and MBP 42kDa), indicative of successful cleavage. (D) Coomassie Gel of fractions from SEC run “mix 4” shows the presence of bands corresponding to the correct molecular weight for RhoA and MBP-Syx_393-792_. Bands in fractions at 8 and 9ml indicate the presence of MBP-Syx_393-792_ and RhoA where bands in fraction 15, 16, and 17ml indicate the presence of RhoA and cleaved MBP-Syx_393-792_ due to background cleavage activity at the TEV site.

**Figure 3.**
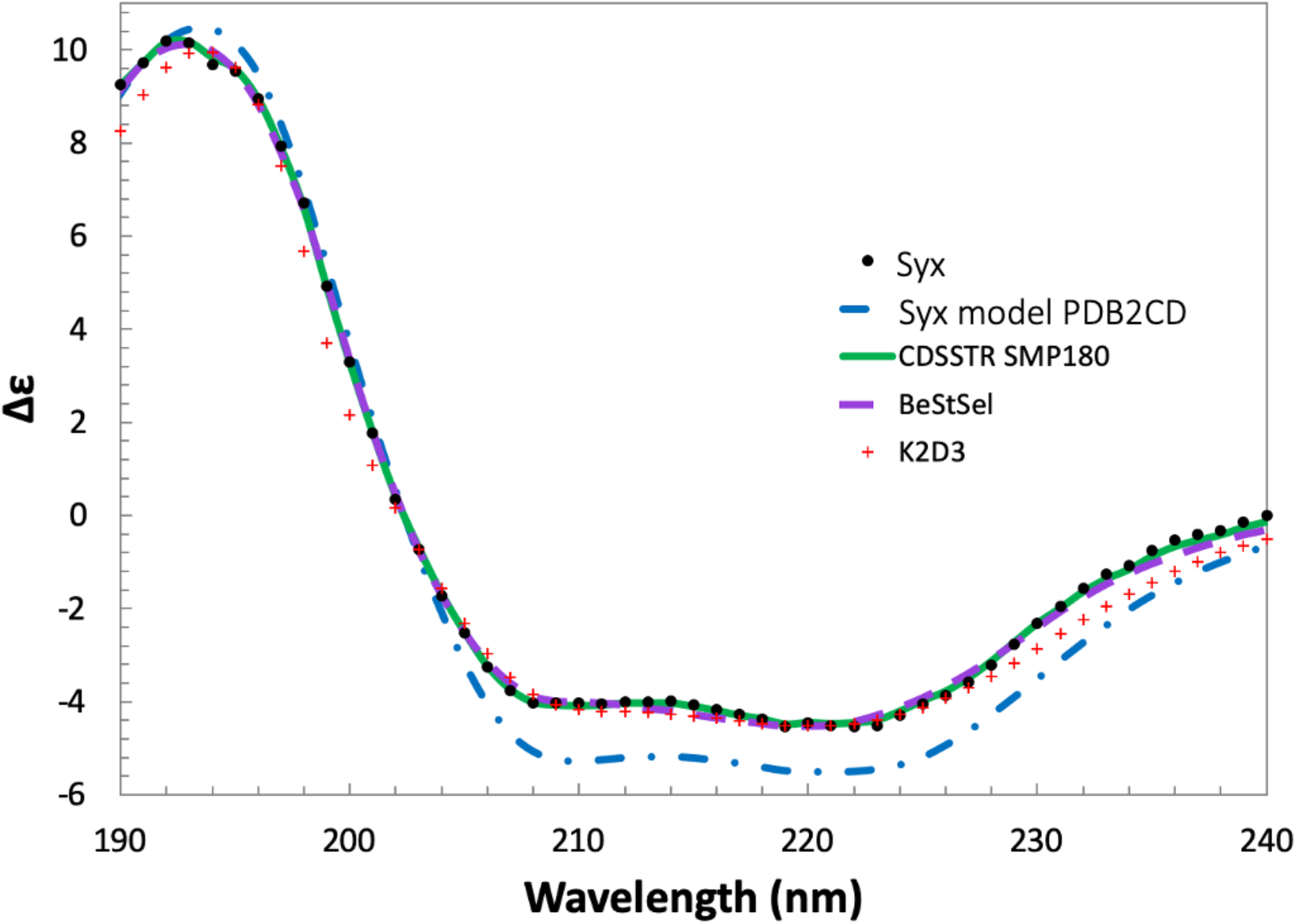
CD spectral analysis of Syx. Experimental spectra is shown as black dots in the plot below. Fits of CDSSTR (SMP180), BeStSel, and K2D3 CD analysis algorithms are shown as purple dashes, green line and red + respectively. The predicted spectra of Syx based on the homology model is shown in blue dashes.

### RhoA – Syx_393-792_ complex formation

Size exclusion chromatography was used to visualize peak shifts that indicate the formation of a protein-protein complex upon mixing MBP-Syx_393-792_ or cleaved Syx_393-792_ with RhoA. The Syx_393-792_-RhoA complex was formed in several different trials to establish conditions which would produce a monodisperse complex suitable for structural studies. RhoGEF DH-PH domains are expected to have the highest affinity for RhoA when it has no nucleotide bound, therefore removal of GDP is likely necessary to drive complex formation with Syx. Most small GTPases are unstable in their apo form and quickly degrade without GDP or a GEF stabilizing them.

This step was complicated by the high binding affinity of GDP with RhoA, which required optimization to find conditions which do not drive aggregation but still effectively remove GDP.

The dashed line in each of the SEC traces in Figure 3A indicates absorbance at 260nm which is the absorbance peak for nucleotides like GDP while the solid line shows the absorbance at 280nm. peaks where the 260nm absorbance signal is greater than 280nm absorbance signal suggest the presence of lingering GDP. These data indicate that GDP appeared to be most effectively removed from RhoA by ammonium sulfate precipitation in the presence of Syx (Figure 3A, second trace from the top, shown in light blue). This condition also shows the largest peak shifted towards lower retention times, suggesting the presence of a complex which was larger than the control trace of Syx alone (Figure 3A, orange trace). This was confirmed by SDS-PAGE analysis showing bands at the correct molecular weights for Syx_393-792_ and RhoA in this fraction (Figure 3C).

### Circular dichroism spectroscopy indicates that E. coli expressed Syx393-792 has characteristic spectra of a folded protein

Circular dichroism (CD) spectroscopy was performed to ascertain if cleaved Syx_393-792_ was folded correctly by comparing the experimentally predicted secondary structure with the secondary structure of known highly homologous crystal structures. Analysis of the CD spectra of Syx predicted an α-helical content of 30–56% and a β-sheet content of 10–24% depending on the algorithm used. CD results were checked by feeding an iTASSER homology model into PDB2CD that resulted in a plausible secondary structure prediction of 47% helix and 11% β-sheet (Table 1)ranjib(57). Therefore, CD indicates the presence of a folded protein (Figure 4) with similar secondary structure characteristics as Syx homologues (Table 1).

**Figure 4.**
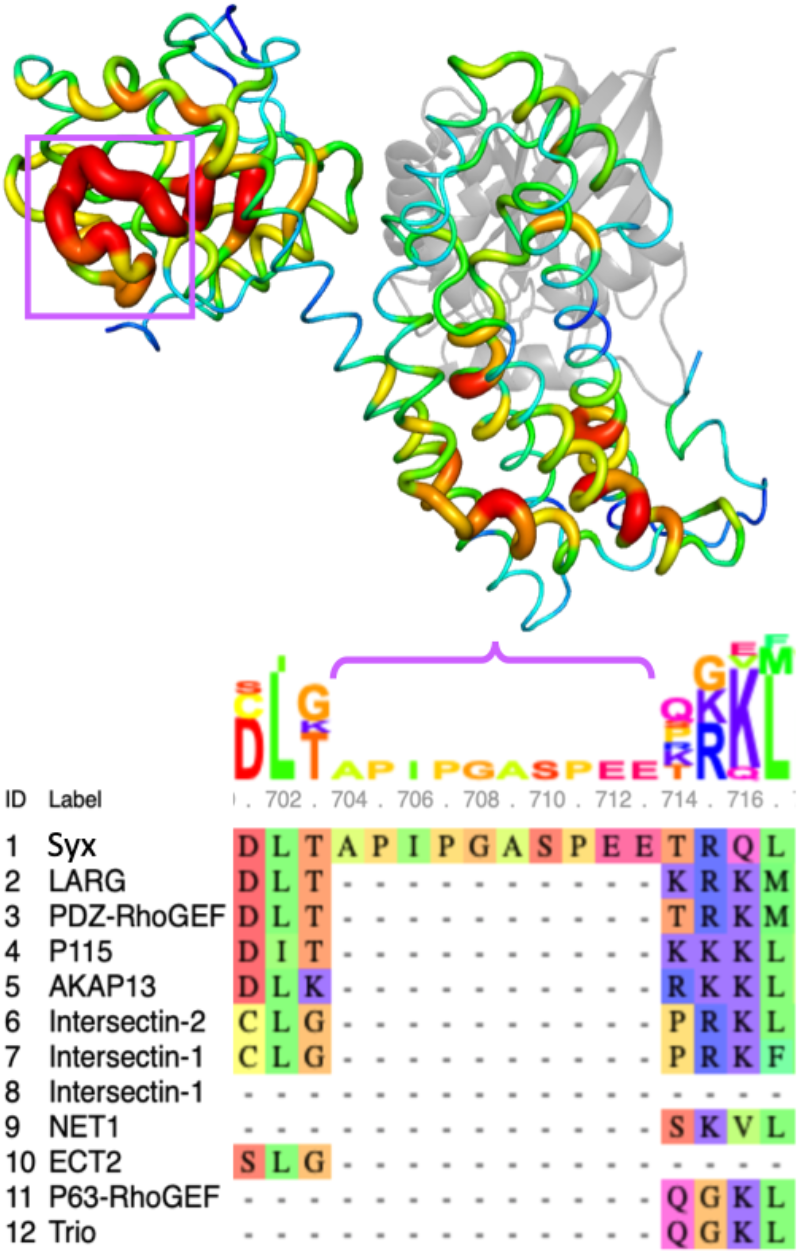
The ratio of non-polar to polar residues that are solvent accessible, as scored by the protein-sol server. Red regions have the highest ratio of non-polar residues while blue patches have the lowest ratio of non-polar to polar residues. The purple bracket indicates the unique hydrophobic loop on the Syx PH domain in the multiple sequence alignment. The analysis reveals a large, membrane facing, non-polar loop on the PH domain which is not conserved in any other homologous RhoGEF structures.

### Biophysical surface analysis

The homology model of Syx was analyzed by several methods to ascertain why size exclusion results suggest the presence of an aggregated protein.

Homology models were analyzed with the protein-sol server that produced protein models scoring the ratio of solvent accessible non-polar residues to polar residues at a given location in the sequence to illustrate relative hydrophobicity of the structure. The analysis also calculated electrostatic potential of the protein surface using a Finite Difference Poisson-Boltzmann (FDPB) method to score the structure such that it can be visualized based on charge distribution (Figure S5)(58).

The analysis revealed a highly un-conserved, hydrophobic loop on the PH domain of Syx that may influence its interaction with the membrane or drive interactions with other hydrophobic domains (Figure 5, indicated in purple). Visualization of charge distribution also showed that the protein was highly polar and had a predominantly positive charge all over the DH domain including the RhoA binding site(59), and a highly negative charge on the PH domain, except for a few membrane facing loops (Figure S5).

**Figure 5.**
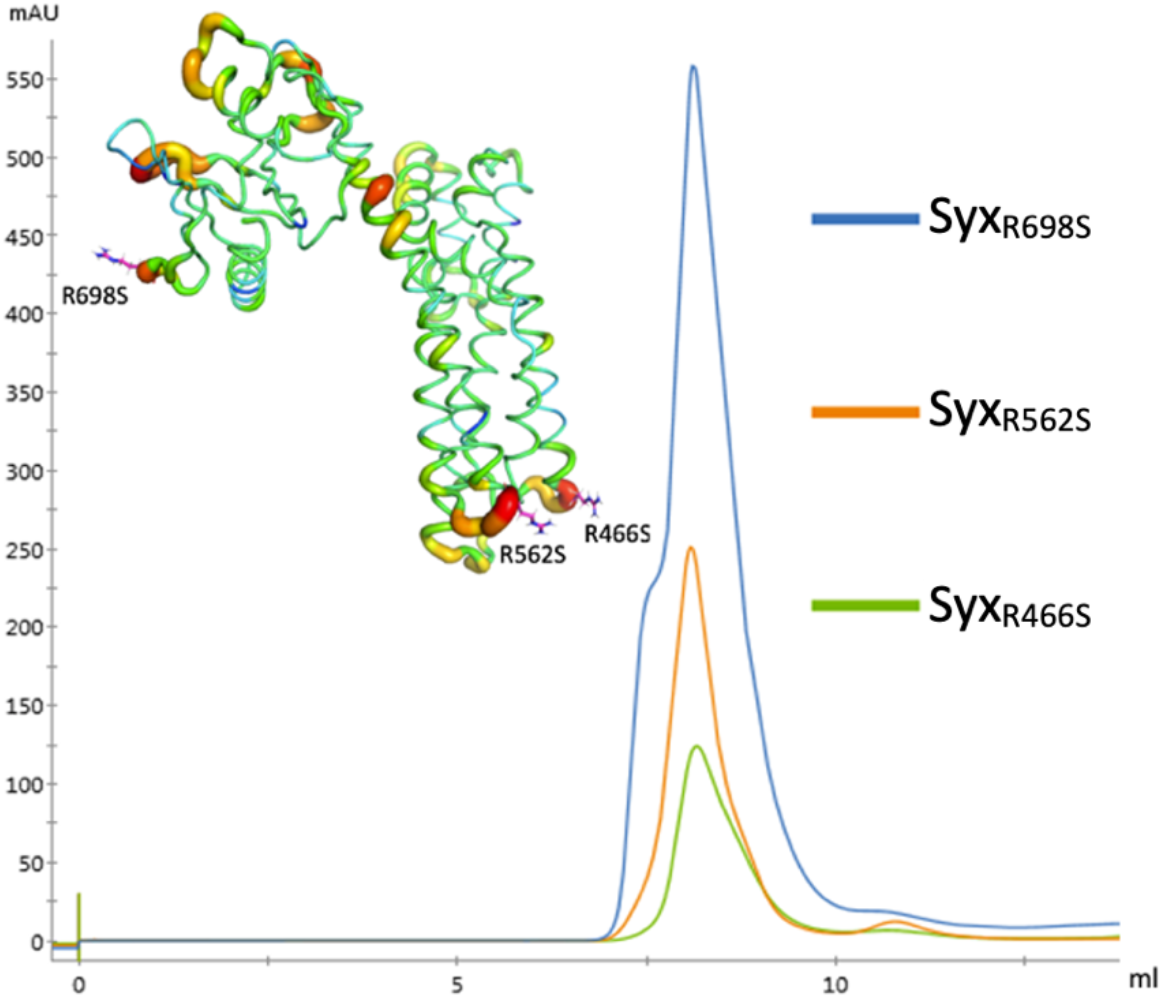
Mutations to surface residues predicted by CamSol web server resulted in SEC peak shifts suggesting that these mutations produced improvements to problematic aggregation characteristics.

### protein engineering

Structurally corrected CamSol web server(46) predictions indicated that several arginine residues were potentially driving aggregation, shown as redder areas on the cartoon (Figure 6). These predictions coincided with several predictions made by Rosetta design for stabilizing mutations. After manual curation of predicted sites, positions R466, R562, and R698 (Figure 6 cartoon) were chosen for site directed mutagenesis. Protein was purified and analyzed identically to Figure 3 MBP-Syx_393-792_ control (orange trace). SEC of mutants revealed several shifts towards higher retention on the column suggesting a reduction in molecular radius of the aggregate (Figure 6 blue, orange, and green trace).

**Figure 6.**
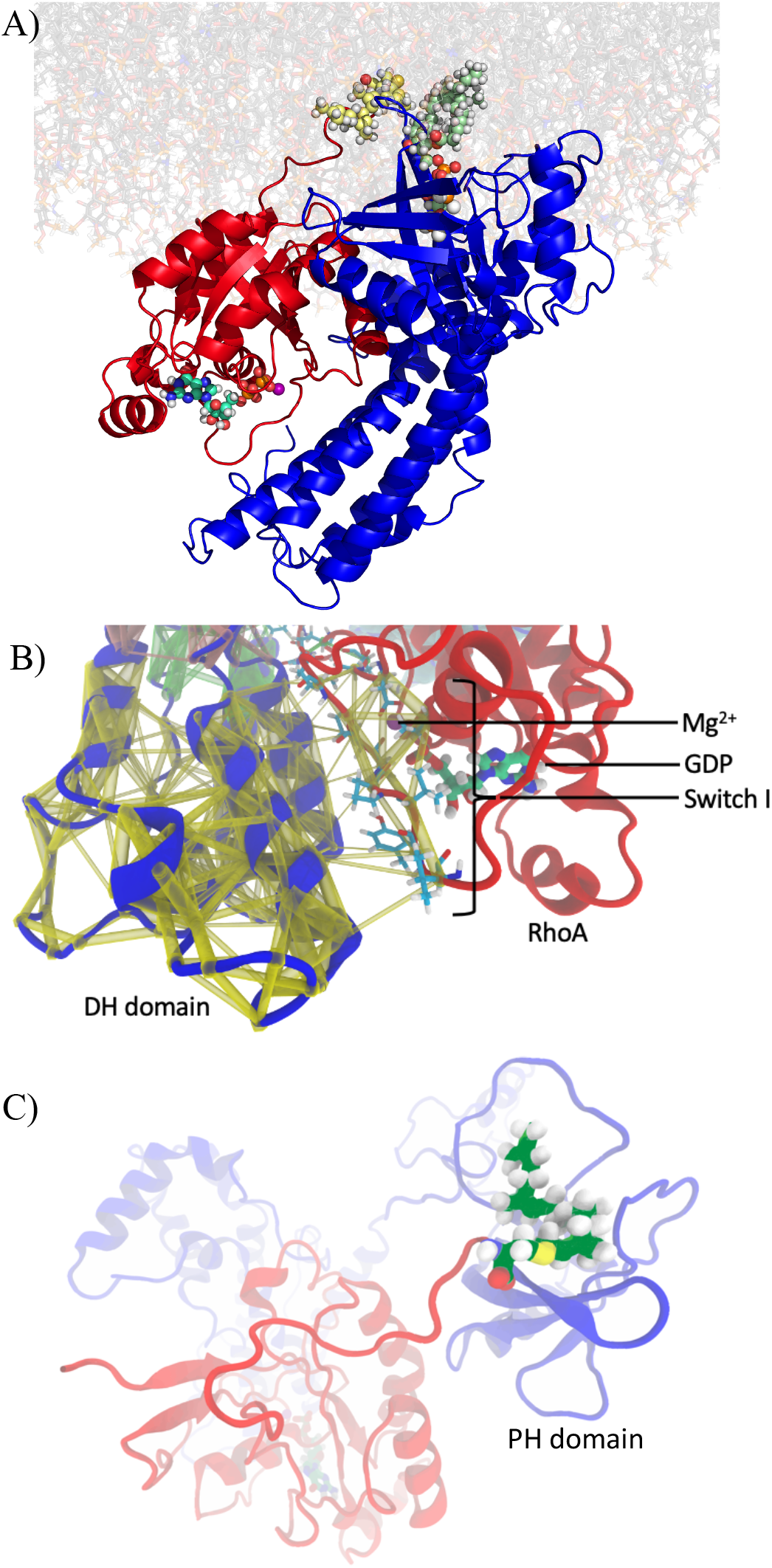
(A) The architecture of the docked Syx-RhoA complex as was simulated, complete with PI(4,5)P_2_ (light green spheres), GDP (teal sticks), and Mg^2+^ (purple sphere) cofactors and post translationally geranylgeranylated Cys-190 (light yellow spheres), simulated on a POPC and PI(4,5)P_2_ bilayer. (B) Dynamic network analysis of the DH domain interacting with RhoA Switch I and Switch II residues (shown as cyan sticks) illustrate how these interactions (yellow tubes) influence the residues responsible for complexing Mg^2+^. (C) shows the geranylgeranyl group (green) conjugated to RhoA (red) residue C190. The geranylgerany group formed transient interactions with the PH domain of Syx (blue) at the interface of the membrane and the cytoplasm during the 1 microsecond all atom mollecular dynamics simmulation.

### Molecular dynamics simulations recapitulate interactions seen in RhoGEF crystal structures and suggest mechanisms of membrane allostery

An all atom molecular dynamics simulation of the full length Syx-RhoA complex (Figure 7A, cropped for visualization) was performed to observe if the protein-lipid interface between Syx and the membrane produced significant structural reorganization of hydrophobic loops, and dynamic network analysis was performed to attempt to detect interactions that might produce allosteric effects on the active site of RhoA.

Dynamic network analysis revealed that these simmulations accurately recapitulated binding interactions that have been observed in other co-crystal structures of Dbl family RhoGEFs with RhoA (Figure 7B)(3, 60). Both the “switch I” and “switch II” region of RhoA were shown to interact with the DH domain of Syx (Figure 7B)(61). Syx interactions with the “switch I” region of RhoA were observed to co-vary with several residues responsible for binding the magnesium co-factor that interacts with the di-phosphate moiety of GDP, including Thr-19, Thr-37, and Asp-59. Displacement of this magnesium is expected to be pivotal for hypothesized mechanisms of guanine exchange(62).

Optimal path analysis to determine the most significant interaction networks between two linked residues suggested that several interactions between the PH domain and RhoA were observed to be capable of linking membrane interacting residues on the PH domain to many key residues in the RhoA switch region such as Thr-37. Residues such as Lys-104 of RhoA were found to propagate interactions from the PH domain all the way to the GDP binding site (Figure S6A), and have the potential to propagate allosteric interactions that could influence GEF activity when the GEF-RhoA complex is in the presence of a membrane.

Additionally, the modeled geranylgeranyl linked cysteine-190 at the C-terminus of RhoA was observed to make several transient interactions with the membrane embedded PH domain of Syx (Figure 7C).

## Discussion

Here we use a combination of protein expression, purification, spectroscopic and modeling techniques to characterize Syx, a partially soluble Dbl RhoGEF protein. Successful expression and purification of Syx was pivotal for all biophysical characterization performed. The exploratory expression of the wild type mouse Syx gene initially displayed low expression and degradation when expressed in *E. coli*. The fact that codon optimization alleviated the formation of degradation products suggests that initial expression difficulties were caused by bacterial ribosomes being unable to complete production of the protein because of stoichiometric restrictions due to the presence of rare or promiscuous codons in the mouse gene. It is also notable that all solved structures of RhoGEFs were fragments of the DH and PH domain, expressed with the help of fusion constructs of either MBP or GST(63–65). We also observed that MBP fusion enabled high level expression and protein solubility dropped dramatically upon TEV cleavage, suggesting that solubility was a culprit of the expression issues with these proteins and that hydrophobic loops might be exposed on the protein surface.

Protocols for production of a monodispersed sample of native Syx protein remain elusive despite numerous exhaustive attempts at construct and buffer optimization, including pH modulation, addition of several detergents, chaotropes, kosmotropes, and charged amino acids (arginine and glutamic acid)(66, 67). Buffer and purification optimization is ongoing with the goal of minimizing aggregation and achieving a polydispersity index (PDI) of <0.2 which is ideal for crystallographic studies.

Low solubility is a major limitation for crystallographic studies which work best with high concentrations of protein. TEV cleavage of the purified MBP fusion protein produced an increasingly turbid suspension at concentrations above 4mg/ml, indicating the formation of large aggregates in solution without the solubility enhancing effects of the fusion tag. This was further corroborated by size exclusion chromatography. We have shown circular dichroism experiments that support the hypothesis that this protein is not forming a misfolded aggregate (Figure 4) and is instead forming non-stoichiometric homo-oligomers. This is further supported by the fact that this protein complex shows RhoA binding activity by SEC (Figure 3) and dot blot (Figure S8). This suggests that disruption of the protein-protein interactions that lead to the formation of the large non-stoichiometric homo-oligomer could result in a protein sample which is suitable for structural studies(68–70).

This hypothesis of non-specific interaction is potentially explained by the results of the protein-sol analysis which shows that this protein contains highly hydrophobic loops which may drive interaction (Figure 5). The analysis also revealed highly charged domains with a large positively charged patch on the DH domain and negative charged regions on the PH domain (Figure S4). These patches may stick to each other without the steric bulk and entropically driven stabilizing effects of disordered regions that are present in the native protein. Protein engineering efforts focused on removing the numerous highly exposed arginine residues and other surface facing residues known to cause nonspecific protein interactions such as methionine, tryptophan, tyrosine, and phenylalanine, presents a promising approach for producing monomeric protein suitable for structural studies that will be suitable for future drug design efforts.

A complex structure of Syx bound to RhoA would be most useful for structurally guided drug design so optimized protocols for forming the complex while maintaining the integrity of the proteins is needed. RhoGEFs have highest affinity for apo RhoA so formation of apo RhoA is essential, however RhoA has picomolar affinity for GDP(71). The active site of RhoA contains a magnesium ion which forms multiple ionic interactions with the phosphate groups of GDP. Strategies to remove the tightly bound GDP require removal of the magnesium, either with excess EDTA, high concentrations of ammonium sulphate, aggressive buffer exchange, or the presence of an active GEF.

Ongoing work seeks to quantify Syx binding affinity to phosphoinositide lipids to characterize contextual protein localization in the cell and potentially establish if allosteric inhibition is possible. Syx is known to bind proteins associated with regulation of cell polarity, cell shape in the CRUMBS polarity complex, as well as stress fiber production and the maintenance of cell-cell junctions(4, 72, 73). In vivo, Syx is bound to the CRUMBS complex via Mupp-1(6, 74, 75). Given the membrane trafficking at this site, it would make sense that the PH domain of this protein would likely bind PI(4,5)P2, the canonical ligand for PH domains(9, 76), because PI(4,5)P2 is also associated with actin cytoskeletal activity modulated by the CRUMBS polarity complex(77, 78). Active RhoA stimulates PIP 5-kinase and subsequent formation of PI(4,5)P2, implicating a positive feedback mechanism where active RhoA results in production of PI(4,5)P2, which recruits Syx to reactivate RhoA as part of a cell’s apical cell polarity program(79, 80). Notably, Syx mediated RhoA activation of kinases is colocalized with angiomotin, another phosphoinositide binding protein with phosphorylation dependent oncogenic potential via the hippo pathway(21, 81). In the future, Large Unilamellar Vesicle (LUV) pull-down assays may answer these questions as well as guide ongoing purification and structural analysis strategies that include the addition of various lipids, membrane mimetics, and protein cofactors that may reduce unwanted interactions and ultimately allow for further characterization.

All atom molecular dynamics simulations revealed a network of interactions that facilitate GEF activity when bound to RhoA. The analysis revealed several interactions between RhoA and the PH domain, mediated by hydrogen bonding networks between positively charged residues and PI(4,5)P2 phospho-inositide headgroups (Figure S5) which may represent a novel mechanism for allostery. This interaction may directly influence the active site of RhoA, as optimal path analysis of the dynamic network showed interactions at the membrane interface of the PH domain share covariance networks that form a link to the active site of RhoA via an optimal path traveling through a lipid headgroup (Figure 6B).

Even more intriguingly, simulations revealed novel interactions between the geranylgeranyl prenyl group ligated to the N-term of RhoA residue 191. These simulated interactions represent a hypothesis for a novel function of the PH domain in membrane associated GEFs. It is conceivable that the PH domain constitutes a lipid binding site situated within the membrane that forms intermolecular interactions with the geranylgeranyl group and helps position RhoA for efficient guanine exchange and alter the binding mode between the DH and PH domains in such a way that it could impinge on the switch regions of RhoA, and help facilitate the opening of the binding site, thus enhancing the release of GDP (Figure S6C). These C-terminal residues and post translational modifications of RhoA are missing from all current GEF-RhoA complex structures. Given that addition of liposomes to activity assays has been shown to enhance activity in other related systems, it is conceivable that this type of interaction is contributing to guanine exchange catalysis. Structural and kinetic studies with GEFs bound to geranylgeranylated RhoA in the presence of a membrane are required to confirm this hypothesis which may be the subject of future work.

Subsequent attempts at generating monodispersed Syx will be made by co-expressing Syx with RhoA in insect cells, as well as purification strategies that take into account the amphiphilic nature of this membrane-associated protein. The modeling efforts included in this paper have provided insights for ongoing future work including mutagenesis efforts to engineer the surface of Syx to replace residues that are likely to contribute to nonspecific protein interactions as well as several provocative insights into the potential effect of membranes on the structure and function of this protein that may support establishment of new druggable sites with further experimental validation.

## Supporting information

Supplemental Rev 2

## Abbreviations

GEF: (guanine exchange factor)
PLEKHG5: (pleckstrin homology and RhoGEF domain containing G5)
Syx: (Synectin associated RhoGEF, alternate name for PLEKHG5)
GEF720: (alternate name for PLEKHG5)
RhoA: (RAS homology ortholog A)
PPIs: (protein-protein interactions)
CD: (circular dichroism)
GBM: (glioblastoma)
PIP: (phospho-inositide phosphate)
SEC: (size exclusion chromatography)
IMAC: (ion metal affinity chromatography)
DH: (dbl homology)
PH: (plekstrin homology)
POPC: (1-palmitoyl-2-oleoyl-sn-glycero-3-phosphocholine)
PI(4,5)P2: (phosphatidylinositol 4,5-bisphosphate or PtdIns[4,5]P2)

## Acknowledgements

This work was supported by the Mayo-ASU Structural Biology Alliance funded by Mayo Clinic and the Biodesign Center for Applied Structural Discovery at Arizona State University. We thank Alex Baker, Daniel Ari Friedman, Megumi Ashley Satkowski, Nattö Satkowski, and John Vant for editing assistance.

