## Supplemental Rev 2 for "Characterization and computational simulation of human Syx, a RhoGEF implicated in glioblastoma"

### Supplemental figures

| C-score | Expected TM-score | Expected RMSD |
| --- | --- | --- |
| 0.19 | 0.74 ± 0.11 | 6.2 ± 3.8 |

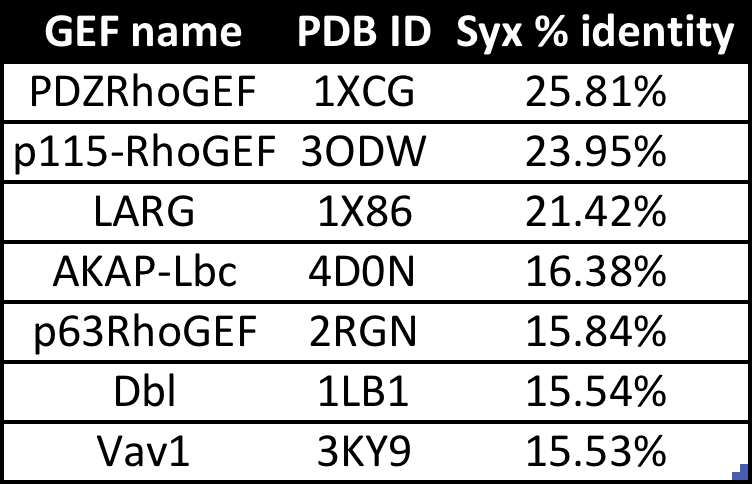

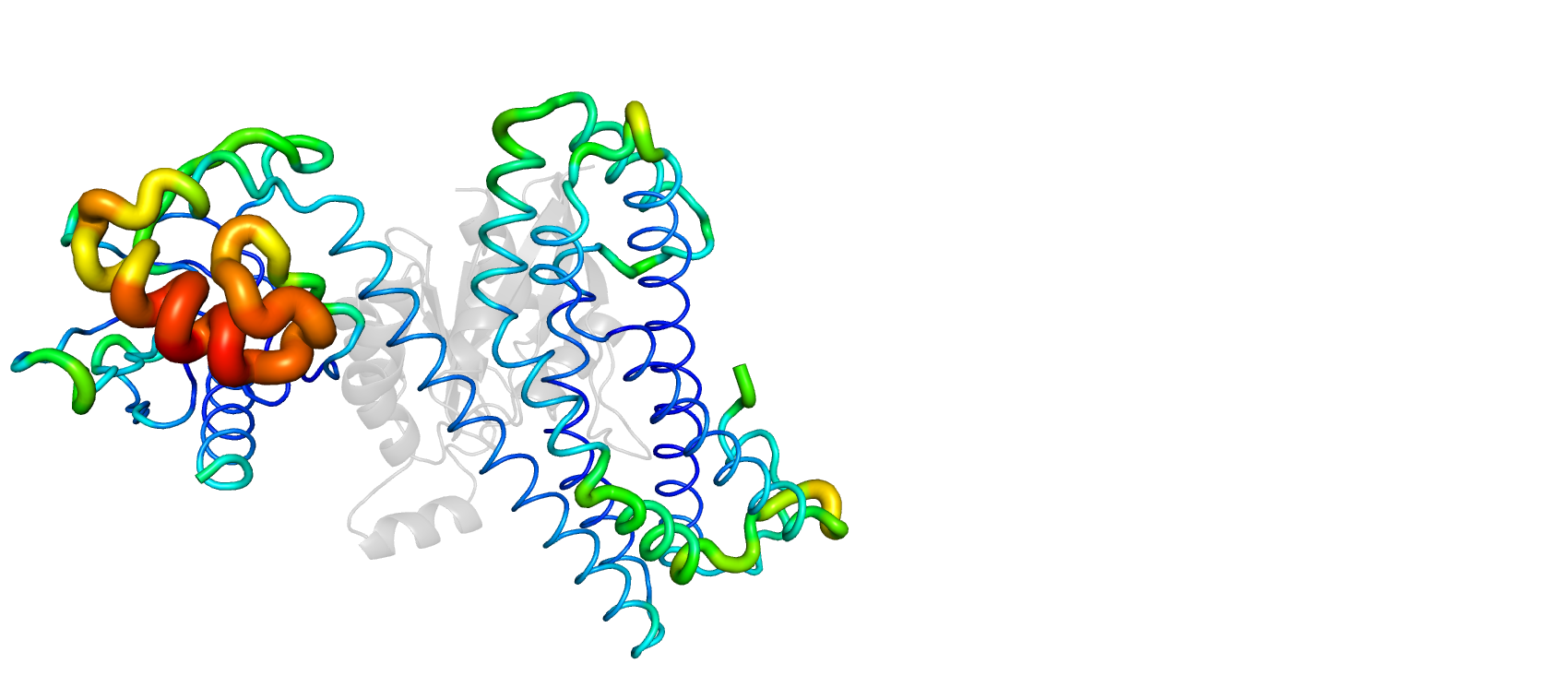

**Figure S1.** (A) Homology modeling of Syx protein produced a homology model. Most key functional regions associated with interactions with RhoA were highly conserved between known PDB structures (shown in blue) but some peripheral loops were not conserved and were therefore poorly modeled as shown in red. (B) Structures in the Protein Data Bank with percent identity to the Syx DH-PH domain of greater than 15% are shown. Three Dbl homology RhoGEFs had the highest percent sequence identity, with PDZRhoGEF sharing 26% sequence identity.

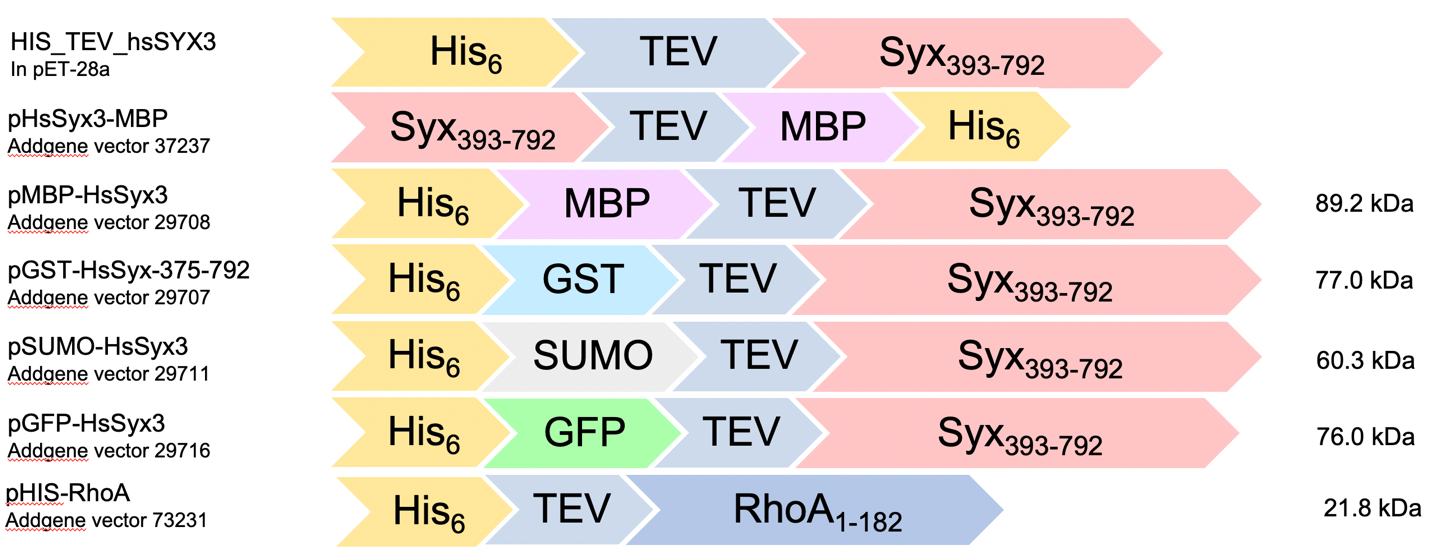
**Figure S2.** All variants of Syx were cloned into addgene constructs based off of the pET series of vectors and were expressed in BL(21) pLysS/I_Q_ *E.coli* cells. Constructs included N-terminal His tagged MBP, GST, SUMO, GFP, and C-terminal MBP with a TEV cleavage sequence for tag removal.

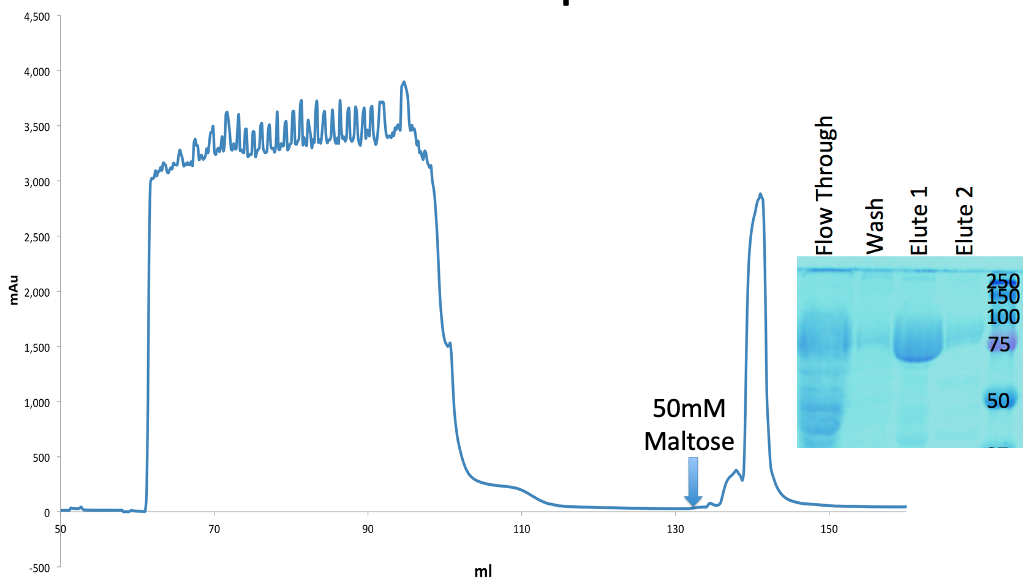

**Figure S3.** Amylose column purification trace of MBP-Syx_375-792_ shown as blue trace produced approximately 90% pure protein as indicated in overlayed Coomassie SDS-PAGE gel image.

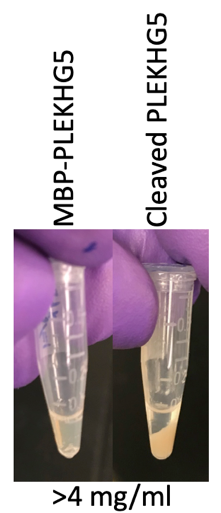

A.

**Figure S4.** Cleavage of MBP from MBP-Syx_393-792_ reduces solubility and results in crashed out protein above 4mg/ml.

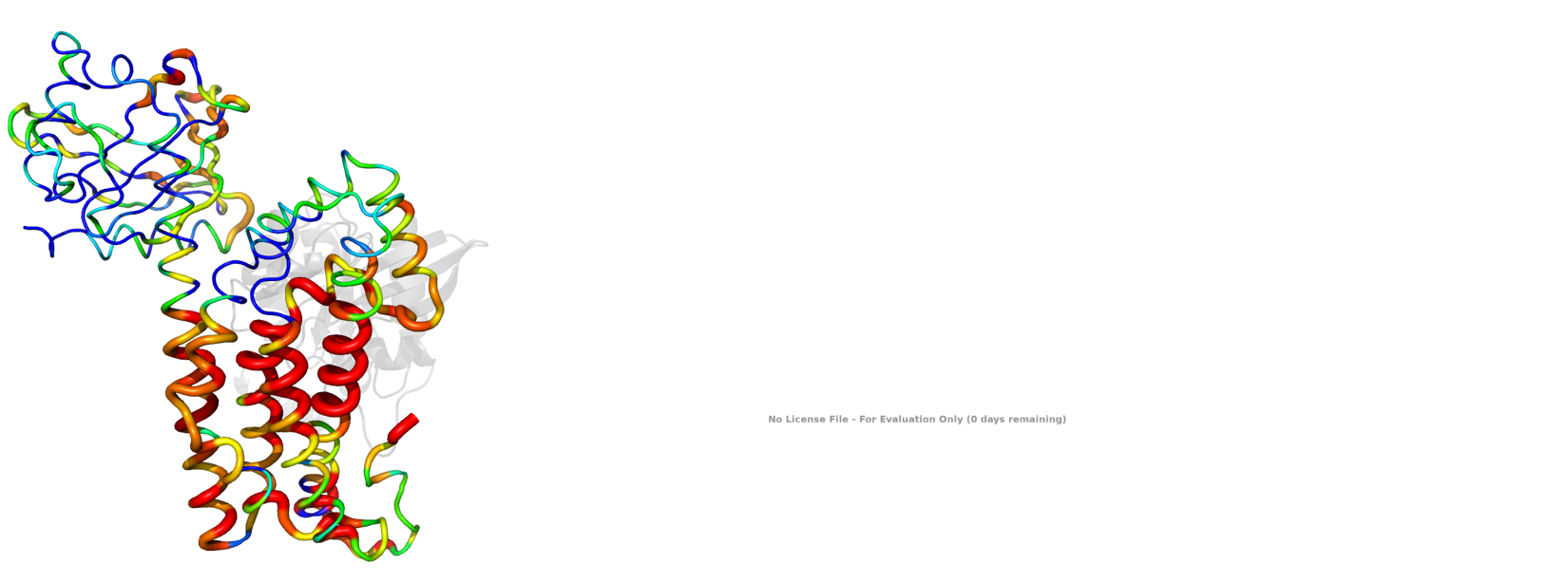

**Figure S5.** B-factor tubes indicate electrostatic potential of the protein surface as calculated by the Finite Difference Poisson-Boltzmann (FDPB) method. Highly positively charged sections colored red and highly negatively charged sections in blue.

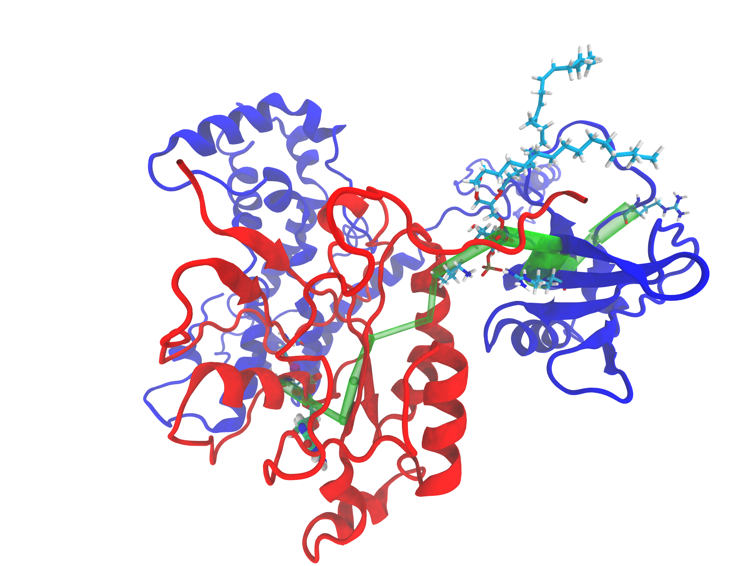

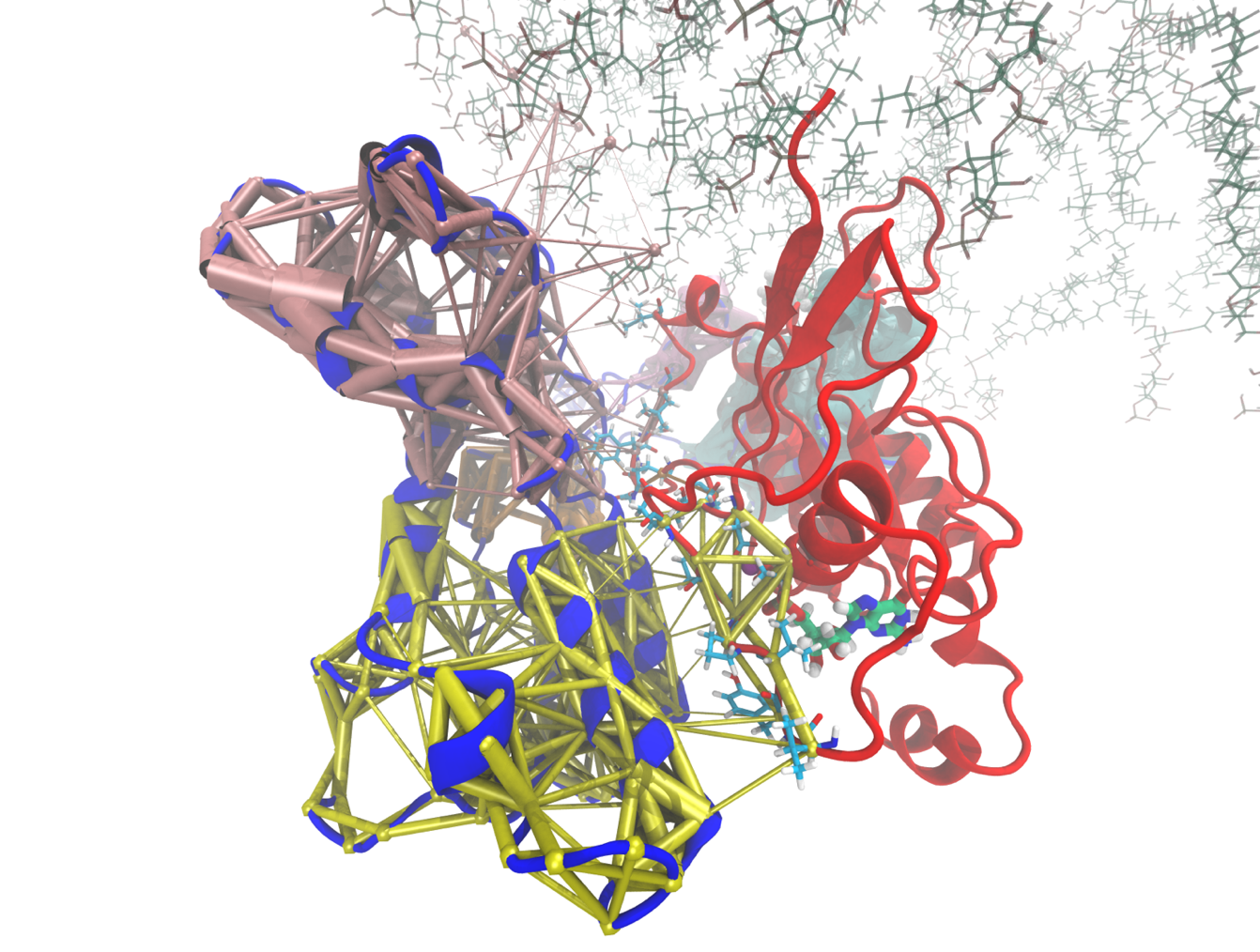

C.

B.

A.

**Figure S6.** (A) Optimal path analysis shown in green shows a direct path from residues at the membrane interface of the Syx PH domain (dark blue cartoon), through a PI(4,5)P lipid (shown as aqua colored sticks), and on to the active site of RhoA (red cartoon). (B) protein-lipid interactions with a modeled lipid membrane (only PA and PI(4,5)P shown to demonstrate membrane orientation), and covariance network of the upper half of the DH domain shows interacting networks of covariance (shown in pink and yellow) propagating from the DH domain of Syx (blue cartoon) into the active site of RhoA (red cartoon). RhoA Switch I residues with significant interactions with the DH domain are shown in aqua, GDP is shown in teal.

**Figure S7.** DLS revealed a polydisperse particle size with a dominant population of particles too large to be monomeric Syx, indicating likely homo-oligomerization or aggregation.

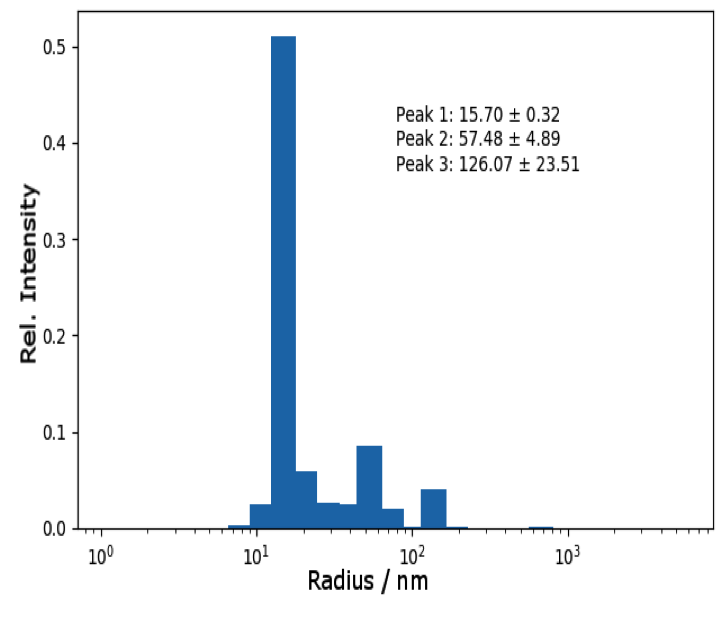

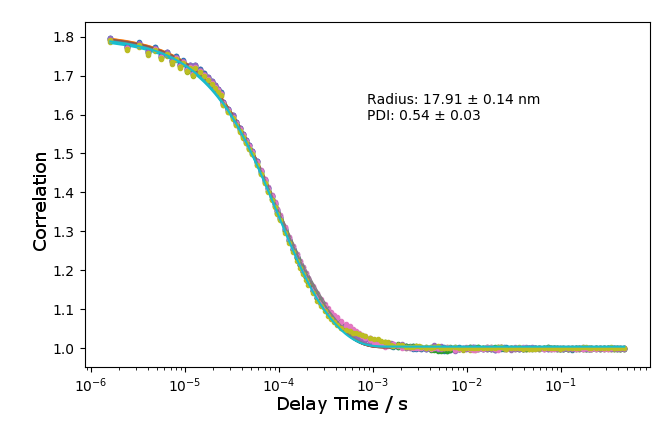
